# *Ganoderma lucidum* alleviates kidney aging by modulating gut microbiota

**DOI:** 10.1101/2024.07.30.605752

**Authors:** Xiaojing Liu, Jiamin Zhao, Jia Liu, Yan Huang, Wei Deng, Luwen Yan, Liao Zhang, Zhihong Liu, Ming Cui, Huiwen Xiao, Xingzhong Liu

## Abstract

**Background:** *Ganoderma lucidum*, a renowned traditional Chinese medicine known for its multiple health benefits, including anti-aging effect, has the ability to influence gut microbiota. However, its specific role in alleviating aging via gut microbiota has not been thoroughly investigated. Here, we explored the impact of *G. lucidum* sporoderm-broken spore powder (*Gl*-SBSP) on kidney aging using naturally aged and radiation-induced premature senescence mouse models.

**Methods:** We analyzed gut microbiota through high-throughput sequencing in naturally aged and radiation-induced premature senescence mice after oral administration of *Gl*-SBSP. Additionally, untargeted metabolomics of the gut microbiome in naturally aged mice was performed using liquid chromatography coupled with tandem mass spectrometry (LC-MS/MS) analyses. Transcriptomics was employed to systematically measure gene expression levels throughout kidney tissues.

**Results:** *Gl*-SBSP effectively mitigated aging-related phenotypic changes and significantly alleviated kidney aging, although outcomes were mixed in the irradiation mouse model. Importantly, *Gl*-SBSP modulated gut bacterial composition, particularly enriching Lachnospiraceae in naturally aged mice and in radiation-induced premature senescence mice with positive prognostic outcomes. Nicotinamide riboside (NR), derived from Lachnospiraceae bacteria, was identified as a potential metabolite that promoted recovery of senescent kidneys and improved renal function, validated through *in vitro* and *in vivo* experiments. Transcriptome analysis revealed that NR regulates kidney steroid metabolism in aging.

**Conclusion:** Our findings suggest that *Gl*-SBSP’s anti-aging effect is mediated by NR derived from the Lachnospiraceae family, influencing kidney steroid metabolism via the gut-kidney axis. These results provide new insights into the mechanisms underlying the anti-aging properties of *Gl*-SBSP.

**Abstract:** *Ganoderma lucidum*, also known as “Linzhi,” is a well-established traditional Chinese medicine renowned for its multiple health benefits and its influence on gut microbiota. In this study, we investigated the anti-aging effect of *G. lucidum* sporoderm-broken spore powder (*Gl*-SBSP) through the gut-kidney axis using naturally aged and radiation-induced premature senescence mouse models. Administration of *Gl*-SBSP not only effectively mitigated aging-related phenotypic changes but also significantly alleviated kidney aging, although outcomes were mixed in the irradiation model. Importantly, *Gl*-SBSP modulated gut bacterial composition, particularly enriching Lachnospiraceae in naturally aged mice and in radiation-induced premature senescence mice with favorable and long-lasting outcomes. Nicotinamide riboside (NR), derived from Lachnospiraceae bacteria, was identified as a potential metabolite that facilitated the rejuvenation of senescent kidneys and improved renal function through both *in vitro* and *in vivo* experiments. Further investigation revealed that NR regulated steroid metabolic process in kidney. Our findings suggest that the anti-aging effect of *Gl*-SBSP is mediated by NR derived from the Lachnospiraceae bacteria, thereby influencing kidney steroid metabolism through the gut-kidney axis. This study provides novel insights into the anti-aging mechanisms of *Ganoderma*.

## Introduction

Aging is an inevitable, long-term process that intensifies over time, leading to the gradual decline or loss of bodily functions. The kidney, being highly sensitive to aging, plays a crucial role in metabolite excretion, essential nutrient re-absorption, and the regulation of metabolic and endocrine functions [1]. As we age, kidney function gradually declines, accompanied by macro and micro-scopic histological changes. These physiological alterations attenuate the kidneys’ ability to repair damage and efficiently excrete waste, increasing the risk of acute and chronic kidney diseases, which in turn accelerates the overall aging process [1,2]. Additionally, with rising life expectancies, age-related conditions such as hypertension [3], diabetes [4] and exposure to nephrotoxic medications are becoming more common [5,6]. These factors contribute to renal impairment, further promoting kidney aging and exacerbating the overall aging process. Currently, the prolonged use of pharmaceuticals and the high costs associated with diagnosing and managing kidney disorders impose a substantial burden on the health and socioeconomic well-being of the elderly. Consequently, finding ways to delay kidney aging has become a pressing concern in clinical settings [7].

Gut microbiota not only help maintain host health and delay aging but also play a critical role in preventing and treating numerous age-related diseases. They achieve this by balancing the gut environment and producing beneficial substances [8–10]. Recently, the interaction between the kidneys and intestines has been termed the “gut-kidney axis” [11]. When the intestinal mucosa is compromised, it loses its ability to prevent harmful bacteria and their byproducts from entering the bloodstream, thereby exacerbating kidney impairment [12,13]. Conversely, kidney injury leads to the accumulation of metabolic waste in the bloodstream, causing additional harm to distant tissues such as the gastrointestinal tract [14,15]. Interventions targeting kidney-related diseases by modulating the gut microbiota have emerged as a novel treatment approach. Strategies such as fecal microbiota transplantation (FMT) [16], probiotics [17,18], antibiotics [19], traditional Chinese medicine (TCM) [20], dietary adjustments and lifestyle changes can be employed to restore the balance of the gut microbiota [8]. The anti-aging effect of gut microbiota have been recognized for over a century [21]. Notably, secondary metabolites from TCM, such as Dendrobium nobile and Polygonati Rhizoma, as well as Klotho, have shown anti-aging effect by protecting the kidneys through modulation of the gut microbiota or microenvironment [22–24].

Traditional Chinese Medicine (TCM) has been extensively accepted and included in the 11th edition of the Global Medical Guidelines by the World Health Organization [25]. Notably, TCM has demonstrated positive impacts on regulating gut microbiota to combat diverse metabolic disorders [26–29]. *Ganoderma lucidum*, renowned as the natural “king of herbs” in TCM, is celebrated for its long-term health benefits and has exhibited a wide range of pharmacological effects, including anti-inflammatory [30], anti-apoptotic [31], antioxidative [32], anti-aging [33], lipid-lowering and hypoglycemic properties [34,35]. Many substances in *G. lucidum* are indigestible and can pass through the digestive tract, interacting with the gut microbiota to maintain host health [36,37]. While the anti-aging and kidney-tonifying effects of *G. lucidum* are well-known in China, the precise mechanisms through which it influences kidney aging and the overall aging processes via modulation of the gut microbiota remain largely unknown.

In this study, we aim to investigate the anti-aging effect of *G. lucidum* sporoderm-broken spore powder (Gl-SBSP), the most commercially prevalent *Ganoderma* product in China, through the gut-kidney axis. We explored this effect and its underlying mechanisms using naturally aged and radiation-induced premature senescence mouse models. Our findings demonstrate that *Gl*-SBSP exhibits significant anti-aging phenotypes by influencing gut microbiota composition and enriching specific bacteria that alleviate kidney aging. Further mechanic investigation indicated that nicotinamide riboside (NR), a key metabolite derived from the enriched bacteria, plays a crucial role in regulating kidney steroid metabolism. This study provides mechanistic insights into the anti-aging properties of *G. lucidum* via the gut-kidney axis.

## 2. Materials and methods

### 2.1 Material and animals

*G. lucidum* sporoderm-broken spore powder (*Gl*-SBSP) was provided by Anhui Biotechnology Co., Ltd (Jiangsu, China). Specific pathogen free (SPF) C57BL/6J female mice, including 8-week-old mice and 12-month-old mice, were obtained from Beijing Huafukang Biotechnology Co., Ltd. Mice were maintained under a 12 h light/dark cycle at the animal experimental center. All experimental procedures were approved by the Animal Ethical Welfare Committee.

### 2.2 Experimental design

Mice were individually labeled and co-housed for each treatment group. Twelve treatments were set up as follows: 1) Control treatment: mice were gavage with saline; 2) *Gl*-SBSP treatment: naturally aged mice were gavage with *Gl*-SBSP at 200 mg/kg/day for 3 months; 3) Irradiation (IR) treatment: 8-week-old mice were gavage with saline after 15 Gy local abdominal irradiation; 4) IR+ *Gl*-SBSP treatment: 8-week-old mice were gavage with aqueous solutions of *Gl*-SBSP at 200 mg/kg/day after 15 Gy local abdominal irradiation; 5) IR-donor treatment: following IR treatment and collecting feces after 7 days; 6) IR+*Gl*-SBSP-donor treatment: following IR+*Gl*-SBSP treatment and collecting feces after 7 days; 7) Fecal Microbiota Transplantation (FMT)-IR treatment: irradiated mice treated with FMT from IR-donor treatment; 8) FMT-IR+*Gl*-SBSP treatment: irradiated mice treated with FMT, in which kidney injury of mice was improved from IR+*Gl*-SBSP treatment; 9) IR+*Gl*-SBSP +broad-spectrum antibiotics (ABX) treatment: following IR+*Gl*-SBSP treatment and treated with ABX in water; 10) IR+*Gl*-SBSP -Responsive treatment: irradiated mice with *Gl*-SBSP replenishment and selected those that were responsive; 11) IR+*Gl*-SBSP -Unresponsive treatment: irradiated mice with *Gl*-SBSP replenishment and selected those that were unresponsive; 12) NR (Macklin, China) treatment: 12-month-old mice were orally gavage with NR for 6 weeks.

### 2.3 Irradiation and dosimetry

A Gammacell® 40 Exactor (Atomic Energy of Ca a Lim, Chalk Rive, Canada) was used for all experiments. Mice were injected intraperitoneally with tribromoethanol to ensure immobility before irradiation. Lead shields were used to protect the upper part of the forelegs and lower part of the hind legs of mice while exposing their kidneys. Then, a single dose of 15 Gy γ-rays local abdominal irradiation was used to induce kidney injury of mice at a rate of 0.88 Gy/min. Mice were sacrificed, and tissues were collected on the 21st day after irradiation. *In vitro* irradiation experiments, HK-2 cells at 80% confluence were treated with a single dose of 8 Gy γ-rays. All control groups were placed in the irradiator for the same period of time but not exposed.

### 2.4 Antibiotics cocktail (ABX)

After irradiation, mice were treated with *Gl*-SBSP by gavage and an antibiotics cocktail in drinking water for 3 weeks to clear the gut microbiota, including 125 mg/L ciprofloxacin (Sigma-Aldrich, Spain), 100 mg/L metronidazole (Sigma-Aldrich, Spain), 50 mg/L vancomycin (Sigma-Aldrich, Spain), 100 U/L streptomycin (Solarbio, China), and 100 U/L penicillin (Solarbio, China).

### 2.5 Fecal microbiota transplantation (FMT)

Adapted from the previously established method. Briefly, the microbiota donor mice were treated with or without *Gl*-SBSP after irradiation, and then the feces were collected. The 0.5 g feces were mixed in 5 mL of 0.9% saline and dissolved for 30 min to obtain a suspension, which was centrifuged to remove sediment. The supernatant was gavage to each recipient mouse.

### 2.6 Pole-climbing Test

The pole-climbing test was performed to evaluate motor coordination ability and bradykinesia, following a previously established method. Each mouse was gently placed on the top of the vertical pole (60 cm from the top, diameter 10 mm, with a rough surface), and the total time taken to reach the bottom was recorded.

### 2.7 Complete blood counts (CBC)

Blood was collected in anticoagulant tubes and analyzed using a blood cell analyzer (DF52, Dymind, China) to evaluate the effects of *Gl*-SBSP on peripheral blood.

### 2.8 Urine, Serum and Tissue Collection

Following labels, urine was collected from each mouse every morning for three consecutive days before euthanasia. Samples were centrifuged at 14 000g for 20 min to remove sediment, and 20 μL of the corresponding three-day urine samples from each mouse were pooled for subsequent analysis. Blood was collected and allowed to clot at room temperature for 2 h. Serum was obtained by centrifugation at 3000 rpm for 15 min at 4 ℃. Mice were euthanized by decapitation, and tissues were collected and divided into two portions. The first portion was fixed in 4% paraformaldehyde (Solarbio, China) for 24 hours and prepared for histopathological examination. The second portion was stored at -80 °C for further analysis.

### 2.9 Histopathology

Fixed tissues were embedded in paraffin, sectioned at 5 μm thickness, and mounted on polylysine-treated slides for hematoxylin-eosin (H&E) (Solarbio, China), Masson (Solarbio, China) and immunofluorescence (IF) staining using standard protocols. Stained sections were examined under a light microscope. For IF staining, slides were deparaffinized, re-hydrated, and subjected to antigen retrieval with citrate buffer. Slides were incubated with 3% H2O2 to quench endogenous peroxidase activity, blocked with 1% BSA, and incubated overnight at 4℃ with anti-Klotho (ab181373) (Abcam, Britain). Subsequently, slides were incubated with fluorochrome-conjugated secondary antibodies (Proteintech Group, USA) in the dark and counterstained with DAPI (Solarbio, China).

### 2.10 Quantification of NAD^+^ levels

Quantification of relative NAD**^+^** levels were performed using the CoenzymeI NAD(H) Content Assay Kit (Solarbio, China) following the manufacturer’s instructions.

### 2.11 Quantitative real-time polymerase chain reaction

Total RNA from tissues and HK-2 cells was isolated using TRIzol (Invitrogen, Carlsbad, CA, USA). Complementary DNA (cDNA) was synthesized using the TRansScript First-Strand cDNA Synthesis SuperMix (TransGen Biotech, Beijing, China) according to the manufacturer’s instructions. qRT-PCR was performed using the PerfectStart Green qPCR SuperMix (TransGen Biotech, Beijing, China) according to the manufacturer’s instructions. The primers used are shown in Supplementary Table S1.

### 2.12 Enzyme-Linked Immunosorbent Assay (ELISA)

Kidney homogenates were prepared by homogenizing 0.1 g of kidney tissues in 300 μL of phosphate buffered saline (PBS). The homogenates were centrifuged at 3000 rpm for 15 min, and the supernatants were utilized to measure Interleukin-1β (IL-1β), transforming growth factor-β (TGF-β), Klotho and Telomerase protein levels using an ELISA kit (mlBio, China) in accordance with the manufacturer’s instructions. Serum and urine samples were centrifuged to remove precipitates, and the levels of interleukin 6 (IL-6) and reactive oxygen species (ROS) in the serum, as well as neutrophil gelatinase-associated lipocalin (NGAL), cystatin-C (Cys-C) and molecule-1 (Kim-1) in the urine, were measured using ELISA kits following the manufacturer’s protocols. Optical density was measured at 450 nm using an Enspire microplate reader (PerkinElmer LLC, USA).

### 2.13 Cell culture

Human renal tubular epithelial HK-2 cells obtained from the American Type Culture Collection (ATCC) and certified to be mycoplasma-free (CRL-2190) were utilized. These cells were cultured in DMEM/F12 medium (Gibco USA) supplemented with 10% (v/v) fetal bovine serum (Gibco, USA) and 1% penicillin/streptomycin (P/S) (Gibco, USA) in an incubator with 5% CO2 at 37 ℃.

### 2.14 Cell proliferation

Cell proliferation was estimated by CCK-8 assay (ABclonal, China). HK-2 cells were plated onto 96-well plates and treated with or without NR (0-1 mg/mL) after irradiation. The CCK-8 reagent was used to measure cell proliferation at 12, 24, or 48 h.

### 2.15 Senescence-associated β-galactosidase staining

HK-2 cells were cultured to 80% confluence in 6-well plates and senescent cells were evaluated by SA-β-Gal Staining after irradiation (Solarbio, China) according to the manufacturer’s instructions. Stained cells were observed under a standard optical microscope.

### 2.16 Bacterial diversity analysis

For feces samples, DNA extraction was performed using the Magnetic Soil and Stool DNA Kit (TianGen, China, Catalog #: DP712). The V3-V4 region of the 16S ribosomal RNA gene was were amplified used specific primer (5′-CCTAYGGGRBGCASCAG-3′) and (5′- GGACTACNNGGGTATCTAAT-3′). Sequencing libraries were generated using the NEB Next® Ultra™ II FS DNA PCR-free Library Prep Kit (New England Biolabs, USA, Catalog #: E7430L) with added indexes. Library quality was assessed using Qubit and real-time PCR for quantification, and the size distribution was evaluated using a bioanalyzer. Quantified libraries were pooled and sequenced on Illumina NovaSeq platform. Paired-end reads were assigned to samples based on their unique barcodes and truncated by cutting off the barcode and primer sequence. The merged paired-end reads were obtained using FLASH (V1.2.11, http://ccb.jhu.edu/software/FLASH/). Quality filtering was performed on the raw tags using fastp (Version 0.23.1) to obtain high-quality clean tags. The clean tags were compared with the reference database (Silva 16S database) using the UCHIME Algorithm (http://www.drive5.com/usearch/manual/uchime_algo.html) to detect and remove chimera sequences. The effective tags were finally obtained, and taxonomic information was annotated using the RDP classifier algorithm (Version 2.2, https://sourceforge.net/projects/rdpclassifier/).

### 2.17 Untargeted metabolomics analysis

Fecal samples from the experimental mice were placed in the EP tubes and resuspended with prechilled 80% methanol by well vortex. Then the samples were melted on ice and whirled for 30 s. After the sonification for 6 min, they were centrifuged at 5,000 rpm, 4°C for 1 min. The supernatant was freeze-dried and dissolved with 10% methanol. UHPLC-MS/MS analyses were performed using a Vanquish UHPLC system (ThermoFisher, Germany) coupled with an Orbitrap Q ExactiveTM HF mass spectrometer (Thermo Fisher, Germany) in Novogene Co., Ltd. (Beijing, China). Samples were injected onto a Hypesil Goldcolumn (100×2.1 mm, 1.9μm) using a 17-min linear gradient at a flow rate of 0.2 mL/min. The eluents for the positive polarity mode were eluent A (0.1% FA in Water) and eluent B (Methanol). The eluents for the negative polarity mode were eluent A (5 mM ammonium acetate, pH 9.0) and eluent B (Methanol).The solvent gradient was set as follows: 2% B, 1.5 min; 2-100% B, 12.0 min; 100% B, 14.0 min; 100-2% B, 14.1 min; 2% B, 17 min. Q ExactiveTM HF mass spectrometer was operated in positive/negative polarity mode with spray voltage of 3.2 kV, capillary temperature of 320°C, sheath gas flow rate of 40 arb and aux gasflow rate of 10 arb. The raw data files generated by UHPLC-MS/MS were processed using the Compound Discoverer 3.1 (CD3.1, ThermoFisher) to perform peak alignment, peak picking, and quantitation for each metabolite. The main parameters were set as follows: retention time tolerance, 0.2 minutes; actual mass tolerance, 5ppm; signal intensity tolerance, 30%; signal/noise ratio, 3; and minimum intensity, et al. After that, peak intensities were normalized to the total spectral intensity. The normalized data was used to predict the molecular formula based on additive ions, molecular ion peaks and fragment ions. And then peaks were matched with the mzCloud (https://www.mzcloud.org/), mzVault and Mass List database to obtain the accurate qualitative and relative quantitative results. Statistical analyses were performed using the statistical software R (R version R-3.4.3), Python (Python 2.7.6 version) and CentOS (CentOS release 6.6). When data were not normally distributed, normal transformations were attempted using of area normalization method. These metabolites were annotated using the KEGG database (https://www.genome.jp/kegg/pathway.html), HMDB database (https://hmdb.ca/metabolites) and LIPID Maps database (http://www.lipidmaps.org/) (Novogene Bioinformatics Technology Co., Ltd.).

### 2.18 Transcriptome sequencing

At the end of the experiment, kidneys were excised from the mice, and RNA was extracted. RNA integrity was assessed using the RNA Nano 6000 Assay Kit of the Bioanalyzer 2100 system (Agilent Technologies, CA, USA). The sequencing library was prepared using the NEBNe UltraTM RNA Library Prep Kit for Illumina (NEB, USA). PCR products were purified (AMPure XP system) and library quality was assessed on the Agilent Bioanalyzer 2100 system. The clustering of the index-coded samples was performed on a cBot Cluster Generation System using TruSeq PE Cluster Kit v3-cBot-HS (Illumia). After cluster generation, the library preparations were sequenced on an Illumina Hiseq platform and 125 bp/150 bp paired-end reads were generated. Raw data (raw reads) of fastq format were firstly processed through in-house perl scripts. In this step, clean data (clean reads) were obtained by removing reads containing adapter, reads containing ploy-N and low quality reads from raw data. At the same time, Q20, Q30 and GC content the clean data were calculated. All the downstream analyses were based on the clean data with high quality. Reference genome and gene model annotation files were downloaded from genome website directly. Index of the reference genome was built using Hisat2 v2.0.5 and paired-end clean reads were aligned to the reference genome using Hisat2 v2.0.5. We selected Hisat2 as the mapping tool for that Hisat2 can generate a database of splice junctions based on the gene model annotation file and thus a better mapping result than other non-splice mapping tools. Feature Counts v1.5.0-p3 was used to count the reads numbers mapped to each gene. And then FPKM of each gene was calculated based on the length of the gene and reads count mapped to this gene. FPKM, expected number of Fragments Per Kilobase of transcript sequence per Millions base pairs sequenced, considers the effects of sequencing depth and gene length for the reads count at the same time, and is currently the most commonly used method for estimating gene expression levels (Novogene Bioinformatics Technology Co., Ltd.).

### 2.19 Statistical analyses

The data are expressed as the mean ± standard error and were analyzed using GraphPad Prism software. The unpaired, two-tailed Student’s *t*-test was employed to assess significant differences between two treatment groups, while one-way ANOVA analysis was used to evaluate differences among groups. *P* <0.05 was considered statistically significant.

## 3. Results

### 3.1. *Gl*-SBSP alleviates kidney aging in naturally aged mice

Naturally aged mice (12 months old) were orally administered *Gl*-SBSP for a duration of 3 months, and various phenotypic parameters were assessed (Fig. 1A). Although *Gl*-SBSP did not affect body weight (Fig. 1B), it significantly attenuated aging-related symptoms such as roughness, shedding, white fur and bradykinesia (Fig. 1C, D). Furthermore, *Gl*-SBSP ameliorated age-disturbed hemogram parameters, including lymphocyte (Lym), neutrophil granulocyte (Neu) and white blood cell (WBC) counts (Fig. 1E, F and Fig. S1A), as well as reduced inflammatory markers in the serum (Fig. 1G). Treatment with *Gl*-SBSP also induced visible changes in the appearance and structure of kidney tissues, including the attenuation of tubular dilatation, epithelial cell shedding and mild fibrosis (Fig. 1H, I and Fig. S1B). *Gl*-SBSP administration decreased levels of inflammatory factors in kidney tissues (Fig. 1J, K), accompanied by a decrease in the expression of pro-aging genes (Fig. 1L) and an increase in anti-aging protein levels in naturally aged mice (Fig. 1M). Moreover, *Gl*-SBSP reduced biomarkers associated with renal function injury (Fig. 1N-1P). Collectively, these results illustrate the anti-aging capacity of *Gl*-SBSP, particularly in attenuating age-related kidney senescence.

**Fig. 1.**
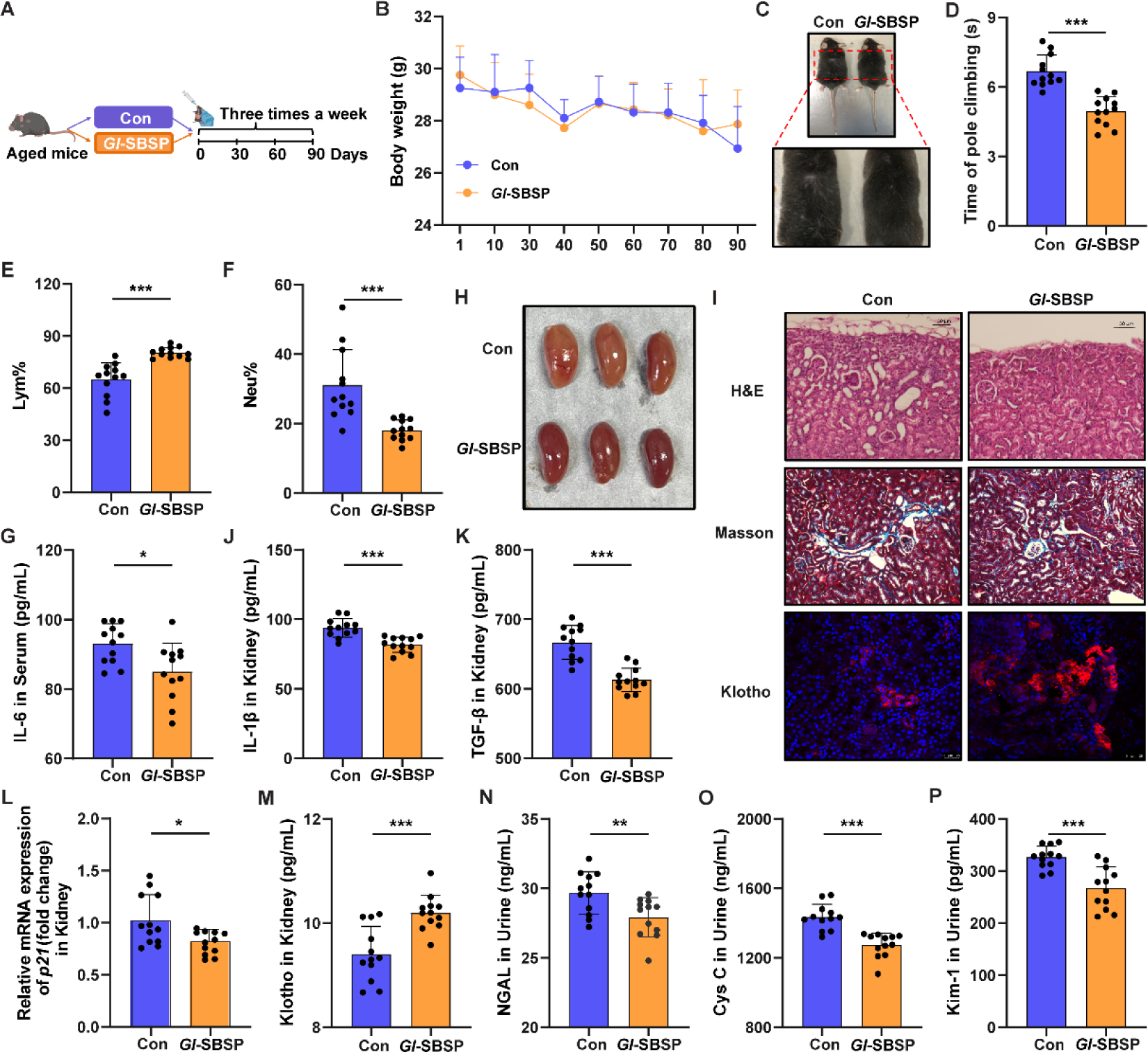
***Gl*-SBSP administration alleviates kidney aging in aged mice.** (A) Schematic diagram of the trial comparing parameters of naturally aged mice with or without *Gl*-SBSP administration after 90 days, with 12 mice for each treatment. (B) Body weight comparison. (C) Body fur appearance of the mice. (D) The time of pole climbing of the mice. (E-F) Percentage of lymphocytes (E) and neutrophils (F) in peripheral blood (PB). (G) Levels of IL-6 in the serum of the mice. (H) Representation of dissected kidney tissues from the mice. (I) Kidney sections stained with H&E (×200 magnification; scale bar: 50 µm), Masson (×100 magnification; scale bar: 50 µm) and IF staining (×200 magnification; scale bar: 25 µm). (J-K) Levels of L-1β (J), and TGF-β (K) in kidney tissues of the mice. (L) Expression levels of *p21* in kidney tissues of the mice. (M) Levels of Klotho in kidney tissues of the mice. (N-P) Levels of NGAL (N), Cys C (O) and Kim-1 (P) in the urine of the mice. * *P* < 0.05, ** *P* < 0.01, *** *P* < 0.001, Student’s *t*-test.

### 3.2. *Gl*-SBSP modulates gut microbiota composition in naturally aged mice

The impact of *Gl*-SBSP on gut microbiota was investigated, revealing a significant increase in the Shannon index for alpha diversity following *Gl*-SBSP administration, despite no visible changes in Observed Features and Chao1 indices (Fig. 2A-D). Weighted non-metric multidimensional scaling (NMDS) and principal coordinate analysis (PCoA) analyses demonstrated a clear separation of microbiota composition between mice with and without *Gl*-SBSP administration (Fig. 2E, F). Both the Firmicutes to Bacteroidetes (F/B) ratio and the frequency of Lachnospiraceae at the family level were enriched after *Gl*-SBSP administration (Fig. 2G and Fig. S2A, B). Heatmap analysis revealed significant alterations in nine genera within the Lachnospiraceae family following *Gl*-SBSP supplementation (Fig. 2H and Fig. S2C-2K). Linear Discriminant Analysis (LDA) effect size assays and T-test analysis further illustrated that the Lachnospiraceae family showed the most significant increase under *Gl*-SBSP exposure (Fig. 2I and Fig. S2L). Additionally, a comprehensive analysis of publicly available databases of human fecal samples from young (20-40 years) and elder (60-80 years) individuals showed a notable reduction in the relative abundance of the Lachnospiraceae family in the elder cohort (Fig. 2J), despite significant variances in gut microbiota composition among young and elder individuals (Fig. 2K) [38]. These data suggest that *Gl*-SBSP supplementation reshapes the gut microbiota in aged mice, particularly by enriching the Lachnospiraceae family. These findings imply that the Lachnospiraceae family might be the target gut bacteria regulated by *Gl*-SBSP for its anti-aging effect.

**Fig. 2.**
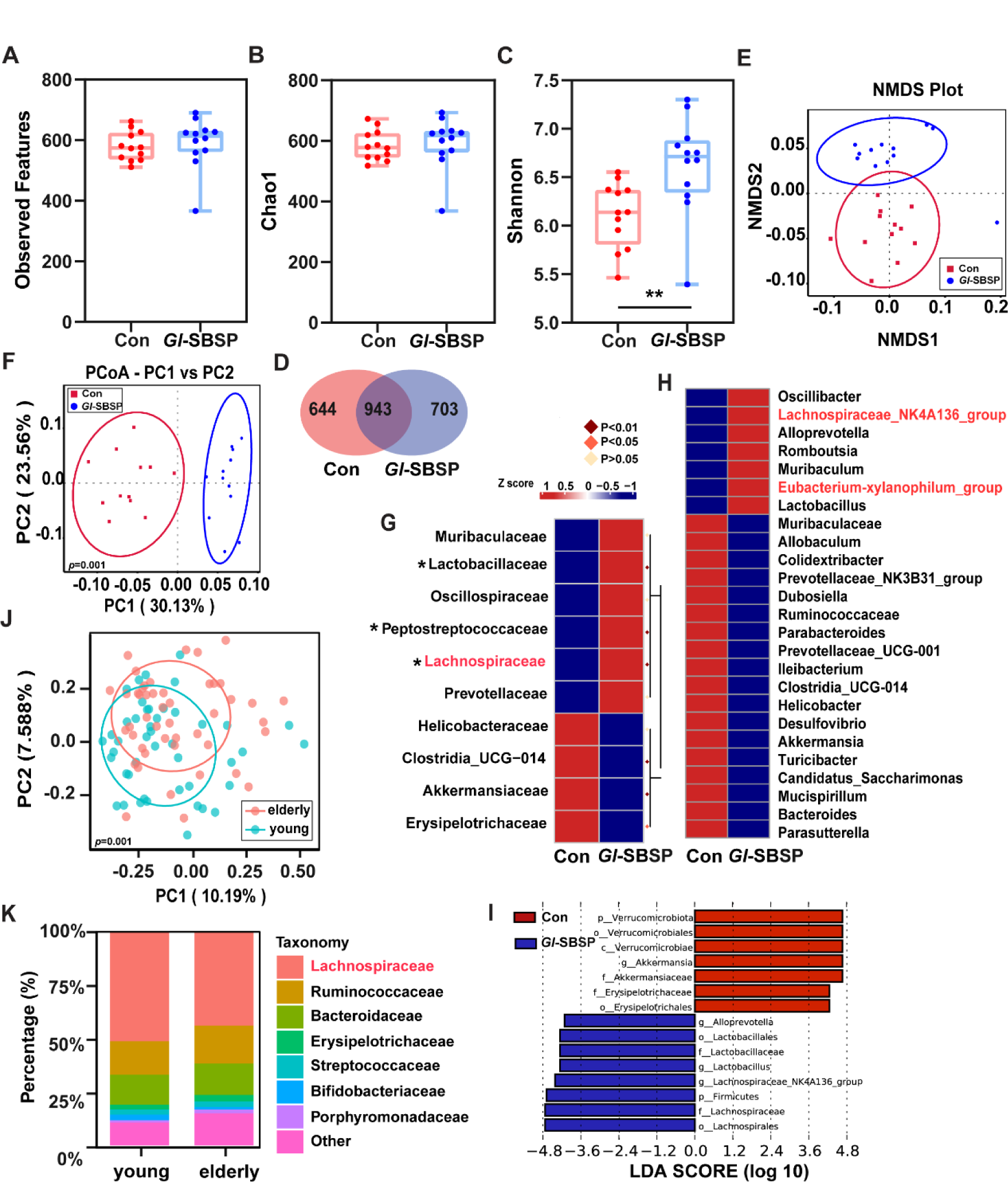
***Gl*-SBSP modulation of gut microbiota affects kidney aging in aged mice, n = 12 per treatment**. (A-C) The Observed Features (A), Chao1 (B), and Shannon (C) diversity indices of intestinal bacteria, derived from OTUs by 16S rRNA high-throughput sequencing. (D) Venn diagrams illustrating the intersections and differences between treatments with and without *Gl*-SBSP administration. (E-F) Shift in intestinal bacterial composition profile illustrated by weighted non-metric multidimensional scaling (NMDS) (E) and principal coordinate analysis (PCoA) (F). (G-H) The alteration of intestinal bacterial patterns at the family (G) and genus (H) level. The heatmap is color-coded based on row Z-scores, with the highest and lowest bacterial levels shown in red and blue, respectively. (I) Linear discriminant analysis (LDA) effect size (LEfSe) showing significant differences in the abundance of gut bacteria and the effect size of each differentially abundant bacterial taxa in the intestines of treatments with and without *Gl*-SBSP administration. (J) PCoA indiagram of the intestinal bacterial composition profile of young and elder individuals from public data. (K) Relative abundance of the top 8 abundant bacteria at the family level in young and elder individuals from public data.* *P* < 0.05, ** *P* < 0.01, Student’s *t*-test.

### 3.3. *Gl*-SBSP protects kidneys against radiation-induced premature senescence of mice

We established a radiation-exposed mouse model to study radiation-induced premature senescence [39,40]. The anti-aging effect of *Gl*-SBSP was investigated using this model (Fig. 3A). *Gl*- -SBSP treatment mitigated radiation-induced premature senescence in the kidneys, though noticeable individual variations were observed. While body weight remained unchanged (Fig. 3B), *Gl*-SBSP increased white blood cell (WBC) count, improved radiation-induced hemocyte myeloid bias in peripheral blood (Fig. 3C, 3D and Fig. S3A), and reduced serum inflammatory levels in irradiated mice (Fig. 3E). Furthermore, *Gl*-SBSP alleviated structural and molecular changes in kidney tissues, evidenced by improved renal atrophy (Figure 3F and Fig. S3B), reduced endothelial cell loss and fibrosis (Figure 3G), lowered levels of IL-β and TGF-β (Figure 3H, I), decreased expression of aging-related genes (Figure 3J, and Fig. S3C), and elevated levels of the anti-aging protein Klotho (Figure 3K). Biomarkers for renal function injury, such as NGAL, Cys C, and Kim-1, were also reduced in the urine following *Gl*-SBSP treatment (Fig. 3L, M and Fig. S3D). *Gl*-SBSP recovered kidney injury and ameliorated renal function without noticeable individual differences within the *Gl*-SBSP treatment group (Fig. S3E-J). Our previous study identified that gut microbiota contributes to the individual differences in response to oral formulations [41]. Thus, we conducted microbiota depletion in irradiated mice using a combination of broad-spectrum antibiotics (ABX) simultaneous with *Gl*-SBSP administration (Fig. 3N). Intriguingly, the individual differences and the protective effect of *Gl*-SBSP were eliminated after ABX treatment. Specifically, ABX introduced a decrease in WBC count and Lym percentage, alongside an increase in neutrophil percentage in irradiated mice treated with *Gl*-SBSP treatment (Fig. 3O-3Q). Levels of IL-6 and reactive oxygen species (ROS) in the serum also showed elevation in ABX-exposed mice (Fig. S3K, L). To further investigate the link between *Gl*-SBSP and gut microbiota, fecal samples were collected from irradiated mice after *Gl*-SBSP administration and transplanted into other irradiated mice (Fig. 3R). Interestingly, the donors showed unconspicuous individual differences and ameliorated premature senescence of hematopoietic system (Fig. 3S-U). These results indicate that the short-term *Gl*-SBSP administration can fight against premature senescence of kidneys, but individual differences in therapeutic efficacy might be influenced by the host’s native gut microbiota.

**Fig. 3.**
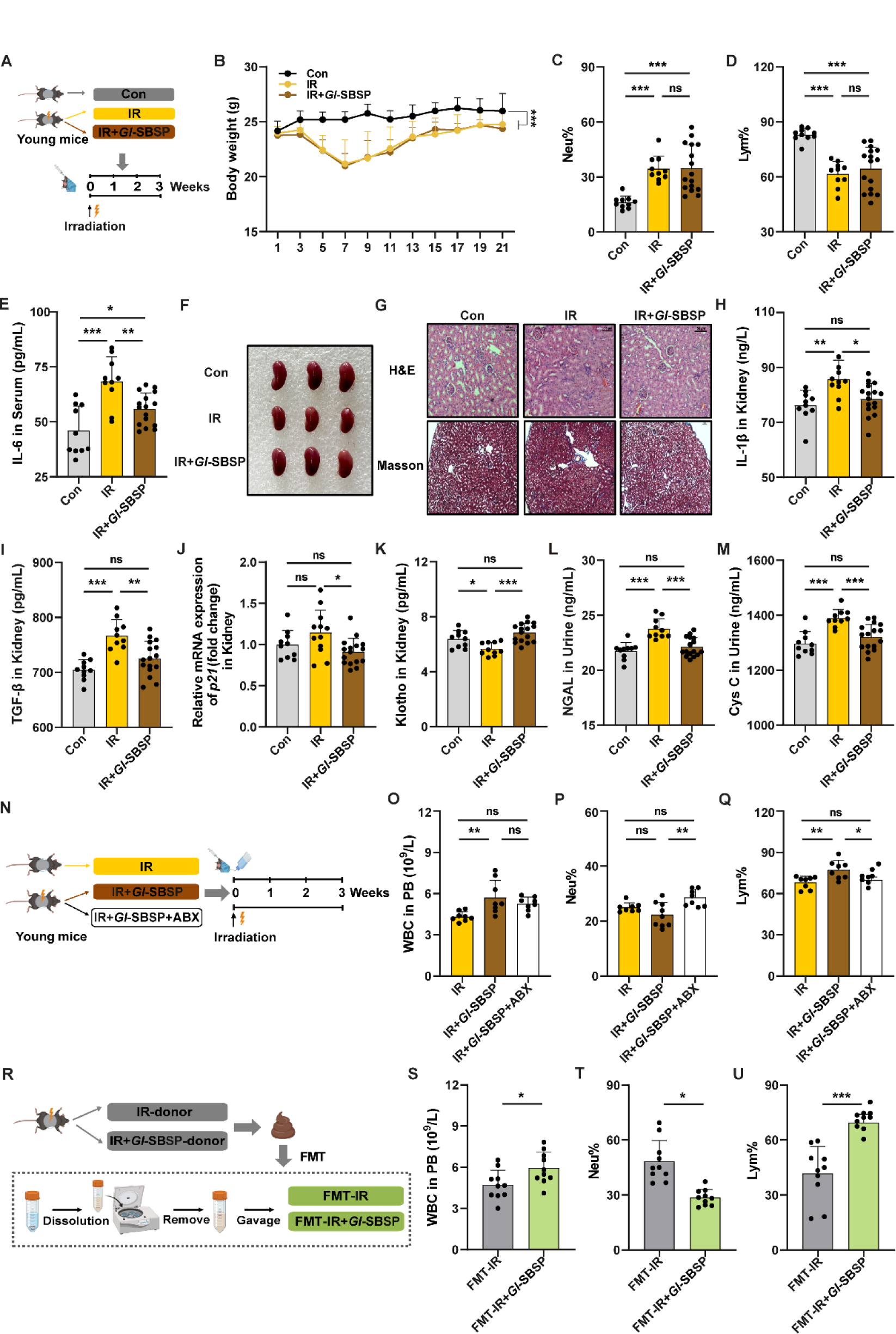
***Gl*-SBSP alleviates radiation-induced kidney aging**. (A) Schematic diagram of the three treatments: mock control, irradiation where the kidney aging mice were exposed to local abdominal irradiation with 15 Gy γ-rays, and *Gl*-SBSP administration in mice after irradiation. All parameters were obtained on the day after three weeks of treatment. (B) The body weight of mice. (C-D) Percentage of neutrophils (C) and lymphocytes (D) in peripheral blood. (E) Levels of IL-6 in the serum of mice. (F) Representative photographs showing dissected kidneys of mice. (G) Representative kidney sections stained with H&E (×200 magnification; scale bar: 50 µm) and Masson (×100 magnification; scale bar: 50 µm) staining. (H-I) Evaluation of IL-1β (H), TGF-β (I) levels in kidney tissues of mice. (J) The expression levels of *p21* in kidney tissues of mice. (K) Levels of Klotho in kidney tissues of mice. (L-M) Levels of NGAL (L) and Cys C (M) in the urine of mice. (N) Schematic diagram illustrating radiation-induced kidney aging in mice exposed to local abdominal irradiation with ABX treatment. (O-Q) Assessment of WBC counts (O), percentage of neutrophils (P) and lymphocytes (Q) in PB. (R) Schematic diagram illustrating radiation-induced kidney aging in mice exposed to local abdominal irradiation with FMT treatment. (S-U) Assessment of white blood cell (WBC) counts (S), percentage of neutrophils (T), and lymphocytes (U) in peripheral blood on day 21 after 15 Gy local abdominal irradiation. * *P* < 0.05, ** *P* < 0.01, *** *P* < 0.001, Student’s *t*-test between each two group, ANOVA among three groups.

### 3.4. The gut microbiota dictates the anti-aging effect of *Gl*-SBSP on radiation-induced premature senescence of kidney

Due to the influence of the native gut microbiome on the anti-aging effect of *Gl*-SBSP, we divided the mice after *Gl*-SBSP administration into “Response” (*Gl*-SBSP-R) and “Un-Response” (*Gl*-SBSP -U) groups based on their physiological indices [42]. Notably, the alpha diversity of the gut microbiota differed among the Control, IR, *Gl*-SBSP-R, and *Gl*-SBSP-U groups (Fig. 4A-D). Weighted NMDS and PCoA analysis showed visible separations of bacterial composition among the groups (Fig. 4E, F). The Firmicutes to Bacteroidetes (F/B) ratio was higher in the *Gl*-SBSP-R group compared to the *Gl*-SBSP-U group (Fig. S4). The abundance of Lachnospiraceae at the family level showed no difference in the gut microbiota between *Gl*-SBSP-R and *Gl*-SBSP-U groups before irradiation (Fig. S5), while increased in *Gl*-SBSP-R group compared to the *Gl*-SBSP-U group after irradiation (Fig. 4G and Fig. S6). Further analysis revealed that the proportion of *Blautia*, *Lachnospiraceae_NK4A136_group*, *Lachnoclostridium* and *Roseburia* within the Lachnospiraceae family were increased in the *Gl*-SBSP-R group (Fig. 4H). T-test and LDA effect size assays demonstrated that Lachnospiraceae exhibited the most significant difference between the *Gl*-SBSP-R and *Gl*-SBSP-U groups (Fig. 4I, J). Taken together with the gut microbial results obtained from naturally aged mice, these findings indicate that the anti-aging effect of *Gl*-SBSP is driven at least partially by the improvement in renal function through the regulation of gut microbiota, particularly Lachnospiraceae.

**Fig. 4.**
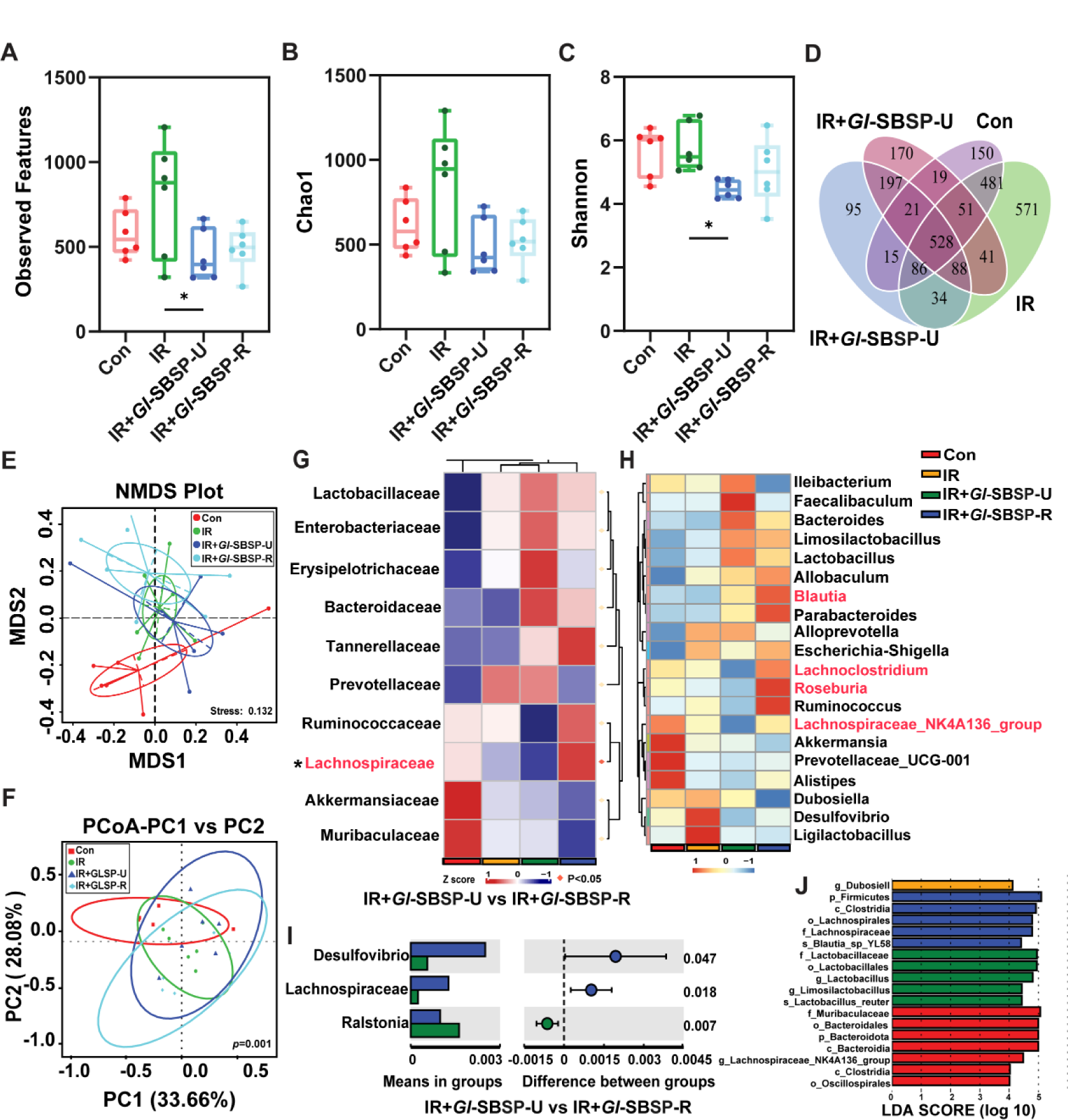
Comparison of intestinal bacterial composition on day 21 after treatments of mock control, irradiation, irradiation following *Gl*-SBSP administration with “Response” (*Gl*-SBSP-R) and “Un-Response” (*Gl*-SBSP-U). (A-C) The Observed Features (A), Chao1 (B) and Shannon (C) diversity index of intestinal bacteria, derived from OTUs 16S rRNA high-throughput sequencing. * *P* < 0.05, ANOVA. (D) The Venn diagrams illustrating the intersections and differences among four treatments. (E-F) Application of NMDS (E) and PCoA (F) to measure the shift in intestinal bacterial composition profile. (G-H) Intestinal bacterial patterns at the family and genus levels using 16S rRNA high-throughput sequencing. The color-coded heatmap based on row Z-scores, with the highest (red) and lowest bacterial levels (blue). (I) T-test analysis for the differences in gut bacterial composition between the Gl-SBSP-U and Gl-SBSP-R. (J) Intestinal abundant bacterial taxa resulting from Linear Discriminant Analysis (LDA) effect size (LEfSe).

### 3.5. Lachnospiraceae-derived NR mediated antiaging

Metabolites are the prime liaisons between the gut microbiota and abenteric organs. To explore the potential anti-aging mechanism of *Gl*-SBSP, untargeted metabolomics analysis and their association with gut bacteria were conducted. It was found that the administration of *Gl*-SBSP remodeled the metabolite profile of the gut microbiota (Fig. 5A, B). A heatmap revealed the difference in the relative abundance of the gut microbiota metabolites, including NR, in the irradiated mice following *Gl*-SBSP replenishment (Fig. 5C). Subsequent conjoint analysis of 16S rRNA high-throughput sequencing and untargeted metabolomics revealed a significant positive correlation between the Lachnospiraceae family and NR, suggesting NR might be a potential key metabolite of Lachnospiraceae (Fig. 5D). Subsequently, we evaluated the anti-aging effect of NR using both cellular and animal models. Our results showed that NR improved the viability of irradiated HK-2 cells (Fig. 5E). NR treatment also reduced the expression of pro-aging genes such as *p53, p21* and *Il-6* (Fig. 5F-H) and decreased the presence of senescent HK-2 cells following irradiation (Fig. 5I). In addition, oral gavage of NR for 6 weeks to naturally aged mice notably improved the roughness, shedding, white furs and bradykinesia, along with a slight increase in body weight (Fig. 5J, K and Fig. S7A). Following NR treatment, there was a significant elevation in NAD^+^ levels in the serum (Fig. 5L), a reduction in Neu percentage, and an elevation in Lym percentage in peripheral blood in naturally aged mice (Fig. 5M and Fig. S7B). Furthermore, the levels of IL-6 and ROS were reduced in the serum (Fig. 5N and Fig. S7C). Within kidney tissues, NR treatment led to an increase in NAD^+^ levels (Fig. 5O), resulting in reduced structural damage and improved renal function (Fig. 5P-R and Fig. S7D, E).

**Fig. 5.**
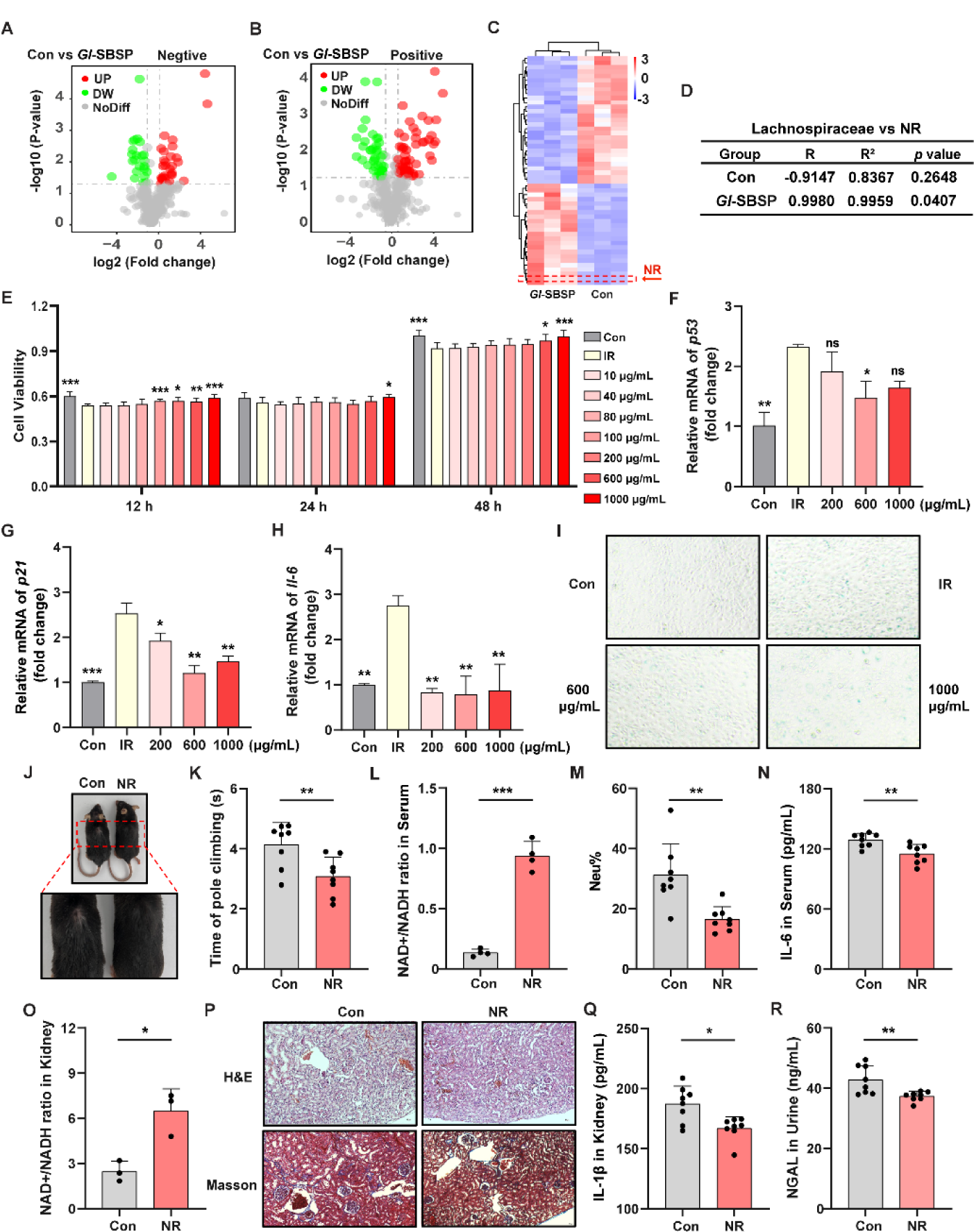
***Gl*-SBSP alleviates kidney aging through elevating NR in naturally aged mice**. (A-B) Volcano plots of different intestinal microorganism metabolites in aged mice. Each point represents a metabolite, with red dots indicating up-regulated metabolites and green dots indicating down-regulated metabolites. (C) Hierarchical cluster analysis of the metabolites in aged mice, with the arrow pointing to NR. (D) Correlation between the bacterial phylum Lachnospiraceae and NR. (E) The proliferation of HK-2 cells were assessed by CCK-8 assays. (F-H) The expression levels of *p53* (F), *p21* (G) and *Il-6* (H) in HK-2 cells exposed to irradiation with 15 Gy γ-rays. (I) Aging cells of HK-2 cells were evaluated by SA-β-Gal Staining after irradiation. (J) Photographs of furs of aged mice. (K) The pole climbing of aged mice. (L) Quantification of relative NAD^+^ levels in the serum of aged mice. (M) Percentage of Neu in PB. (N) Levels of IL-6 in the serum of aged mice. (O) Quantification of relative NAD^+^ levels in kindney tissue. (P) Kidney sections stained with H&E (×100 magnification; scale bar: 20 µm) and Masson (×100 magnification; scale bar: 20 µm). (Q) Levels of L-1β in kidney tissues of aged mice. (R) Levels of NGAL in the urine of aged mice. * *P* < 0.05, ** *P* < 0.01, *** *P* < 0.001, Student’s *t*-test.

### 3.6. NR alleviates kidney aging via regulating lipid metabolism

Following the gut-kidney axis, the gene expression profile of kidney tissues was analyzed after NR gavage in naturally aged mice, with saline as the control. KEGG and GO analyses indicated that NR treatment inhibited the steroid synthesis pathway (KEGG: mmu00100), steroid metabolic and lipid biosynthetic process (Fig. 6A, B), but activated the fatty acid degradation pathway (KEGG: mmu00071), PPAR signaling pathway (KEGG: mmu03320), ATP metabolic process, fatty acid β-oxidation, and fatty acid metabolic process (Fig. S8A, B). We also identified the significantly regulated genes *Msmo1*, *Hsd17b7*, *Ehhadh* and *Hmgcs2*, which are related to steroid biosynthesis, steroid metabolic process, and fatty acid degradation pathways, respectively (Fig. 6C-F). We further validated the expression of these genes with an expanded sample size, which revealed that NR indeed downregulated the expression of *Hsd17b7* and *Ehhadh*, while upregulated *Hmgcs2* in kidney tissues with NR treatment (Fig. 6G-I).

**Fig. 6.**
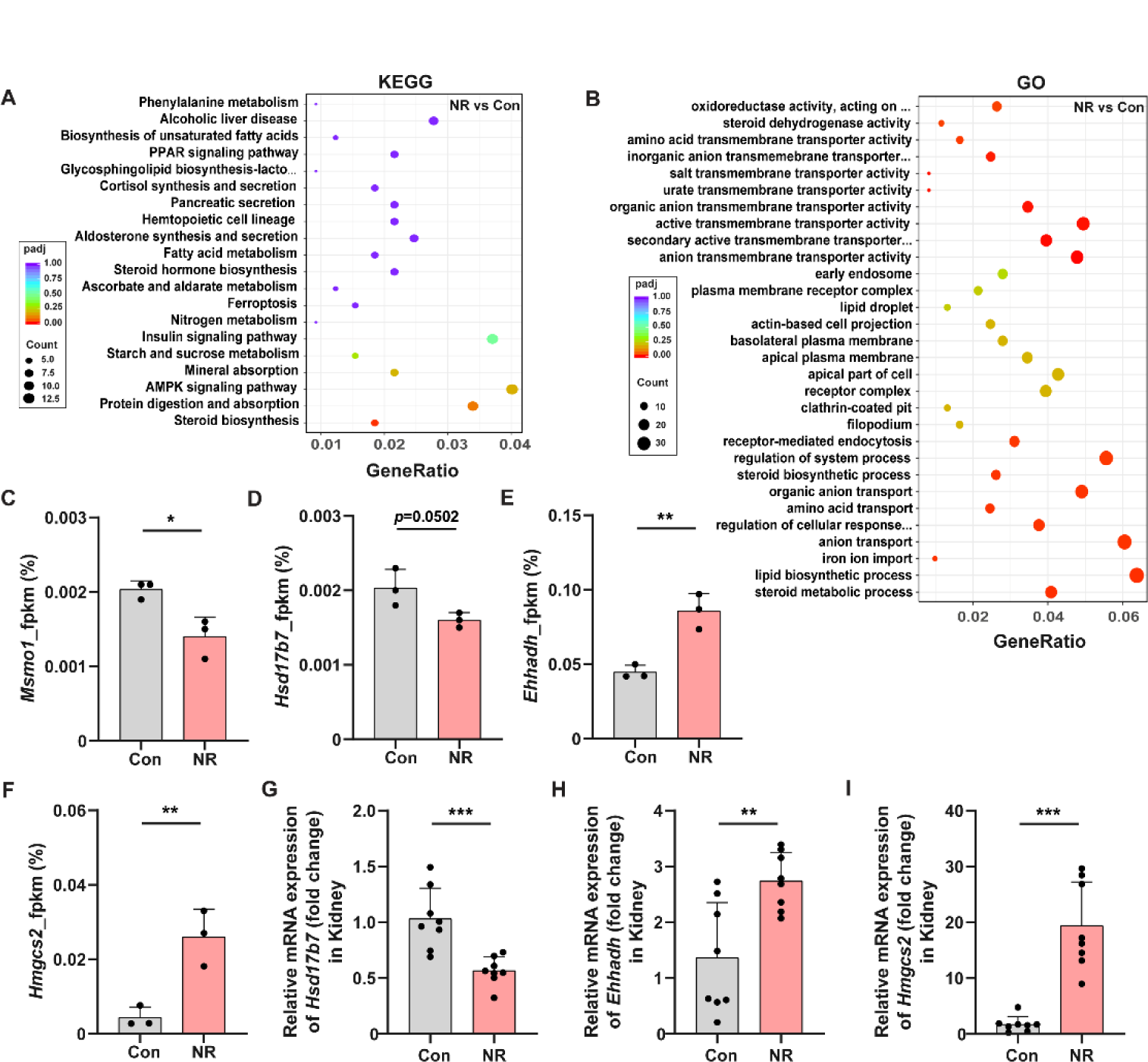
**NR relieved kidney aging via via regulating lipid metabolism**. (A) KEGG pathway analysis of the downregulation of differential metabolites in two groups. (B) GO pathway analysis of the downregulation of enriched metabolites in two groups. (C-F) FPKM of *Msmo1*(C), *Hsd17b7* (D), *Ehhadh* (E), and *Hmgcs2* (F) of aged mice. (G-I) The expression levels of *Hsd17b7* (G), *Ehhadh* (H) and *Hmgcs2* (I) of aged mice. * *P* < 0.05, ** *P* < 0.01, *** *P* < 0.001, Student’s *t*-test. Table S1 Primer sequences for qPCR analysis (qRT-PCR)

## Discussion

*G. lucidum*, cherished for millennia as a miraculous tonic, is renowned for its abilities to tonify the body, promote vitality, and slow the aging process. Recognized by Western societies as nutraceuticals, *G. lucidum* is consumed as health products, nutritional supplements, cosmetics and functional foods [43]. Preclinical and clinical trials have validated the benefits of *G. lucidum* in addressing aging and age-related ailments [44,45]. Recent research has highlighted its role in gut microbiota and intestinal barrier function, mediating the inhibition of obesity [46,47], diabetes [48] and related metabolic disorders [49,50]. Crucial beneficial symbiotic bacteria, such as Lactobacillus, Lachnospiraceae NK4A136, and Ruminococcaceae UGG-014, have been enhanced by *G. lucidum* [51–53]. In this study, we found that *Gl*-SBSP countered kidney aging in both naturally aged and radiation-induced premature senescence mouse models, enriching the gut bacteria of Lachnospiraceae. The anti-aging effect is supported by previous study [54] and the public datasets indicating a decline in gut Lachnospiraceae abundance with aging through comparisons of gut microbiota between young and aged individuals. Additionally, we identified the key NR, potentially derived from Lachnospiraceae, which regulates steroid metabolic processes in the kidney. Our findings provide new insights into *Gl*-SBSP ’s anti-aging effect and mechanisms via the gut-kidney axis.

One of the intriguing phenomena in this study is the different responses to the short-term *Gl*-SBSP treatment among the individuals in the radiation-induced premature aging mice model, as opposed to the natural aging model mice. The *Gl*-SBSP-responsive group of mice significantly enriched Lachnospiraceae bacteria compared to the *Gl*-SBSP-unresponsive group, despite the initial mice being identical and randomly selected. Further investigation through antibiotic (ABX) and fecal microbiota transplantation (FMT) experiments provide compelling evidence for the factors driving an individual’s reaction to *Gl*-SBSP. Such individual differences in response to dietary interventions have been noted in other studies as well. For example, genetically identical twins have shown different responses in blood glucose, insulin, and lipid levels to the same meals [42,55,56]. In pediatric cases of inflammatory bowel syndrome (IBS), children who positively responded to a low fermentable oligosaccharides, disaccharides, monosaccharides and polyols (FODMAP) diet tended to harbor high levels of Bacteroidaceae, Erysipilotrichaceae and Clostridiales species in their gastrointestinal tract. In contrast, the non-responders often harbored high level of *Turicibacter* [57]. Traditional Chinese medical science emphasizes personalized medical treatment that generally depends on an individual’s constitution (physiological condition). The different responses of individuals to various treatments may be affected by the individual constitution, which remains a complex interplay of factors that are not yet fully understood.

Bioactive metabolites are the key mechanism through which the gut microbiota in the gut-X axis affects distal organs via the bloodstream, maintaining the health and homeostasis of these organs [58–61]. Short-chain fatty acids have been widely recognized as active metabolites of gut microbiota, especially Lachnospiraceae [62]. However, comprehensive microbiome and metabolomics analysis indicated that short-chain fatty acids were not among the top 20 metabolites detected in the feces of naturally aging mice. Notably, NR is the most enriched metabolite after administration of *Gl*-SBSP and showed a significant positive correlation with Lachnospiraceae. The comparative assessments have shown that the gut microbiota structure, particularly the enrichment of Lachnospiraceae in centenarians, elderly, and aged mice with NR intervention is notable (Fig. S9) [63–65]. Significant positive correlations between NR, including A2, and both Lachnospiraceae UCG-010 and Lachnospiraceae UCG-006 have been observed (Fig. S2). Furthermore, a homologous search of the Lachnospiraceae family genome also verified the existence of genes related to NR synthesis within the genome (Fig. S10). Although those results highlight the association between Lachnospiraceae and NR and their antiaging effect, we could not provide direct evidences that Lachnospiraceae produces NR from transcriptome and metabolism data due to the failure to isolate and culture those bacteria.

Accumulating evidence confirms that NR requires interaction with the gut microbiota in the small intestine for its conversion into nicotinic acid or nicotinamide in distal organs. Depletion of the gut microbiota in sterile mice or through antibiotic treatment significantly hinders the conversion of NR, leading to a notable decrease in its content across various organs [66–69]. In this study, we demonstrate that NR supplementation elevates NAD^+^ levels in the blood and kidneys of aged mice, thereby slowing renal and overall aging by modulating lipid metabolism pathways in kidney tissue. This effect is attributed to the well-established function of NR as a crucial precursor for NAD^+^ synthesis. NR can enhance tissue NAD^+^ content [70–73], promoting energy expenditure, thermogenesis, inhibiting lipid accumulation and cholesterol levels, and even extending the lifespan of yeast, nematodes, and mice [74–76]. Additionally, NR is ability to repair DNA damage, suppress cell apoptosis and aging by down-regulating the expression of pro-inflammatory cytokines, resulting in promising outcomes in age-related diseases [66,77,78]. Although encouraging results have been achieved in animal models, the therapeutic evaluation of NR in human trials remains limited. Further research is needed to deepen our understanding of NR’s mechanisms of action and its potential as a therapeutic option in clinical settings.

## Conclusion

In summary, *G. lucidum* has a significant anti-aging effect by regulating gut microbiota and their metabolites, especially enriching Lachnospiraceae bacteria and NR to reprogram the gene expression profile especially lipid metabolism processes in kidney aging. It also highlights individual differences in gut microbiota responses to dietary interventions. Those findings unveil mechanistic insights into *Gl*-SBSP as a dietary intervention for the anti-aging as well as for prevention and management of age-related kidney diseases through gut-kidney axis. This provides a pivotal reference for public consumers and clinical settings to promote kidney health with *G. lucidum* products.

## Supplementary data

### Correlation analysis between bacterial taxonomy and metabolites

The correlation between the Lachnospiraceae family and NR was estimated by Pearson’s correlation coefficient analysis. The significant correction was performed with the criteria of \**p* <0.05.

### Homologous search of NR synthesis genes in Lachnospiraceae genome

Genomic data retrieval was conducted in the NCBI database (https://www.ncbi.nlm.nih.gov/) using the keywords Blautia, Eubacterium, Lachnoclostridium, and Roseburia (detected in this study) in the Lachnospiraceae family on June 20, 2024. Reference genomes of each species were selected for subsequent analysis, resulting in a total of 64 genomic data obtained, including 26 from the Blautia genus, 16 from the Eubacterium genus, 11 from the Lachnoclostridium genus, and 11 from the Roseburia genus (Table S1). All protein sequences from the 64 genomes were merged to construct a Lachnospiraceae gene database using DIAMOND software. Additionally, the protein sequences of the five genes associated with nicotinamide ribose synthesis (pncB, nadD, nadR, nadE, and ushA) were retrieved from the NCBI database for *Escherichia coli*. Subsequently, using the DIAMOND software, homology searches were conducted between the pncB, nadD, nadR, nadE, and ushA genes of *Escherichia coli* and the established Lachnospiraceae gene database. Python 3.8 was utilized to calculate the results of the homology searches and a heatmap was generated to display the findings.

---

**Table S1.**
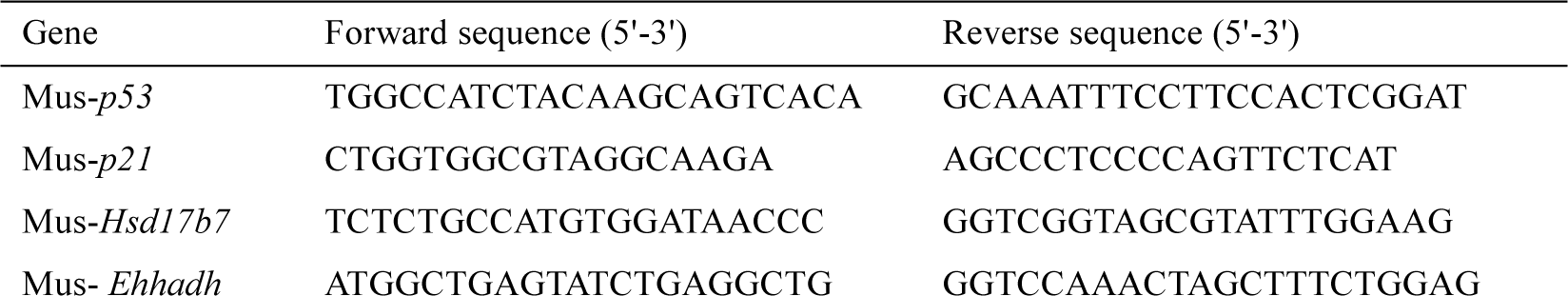
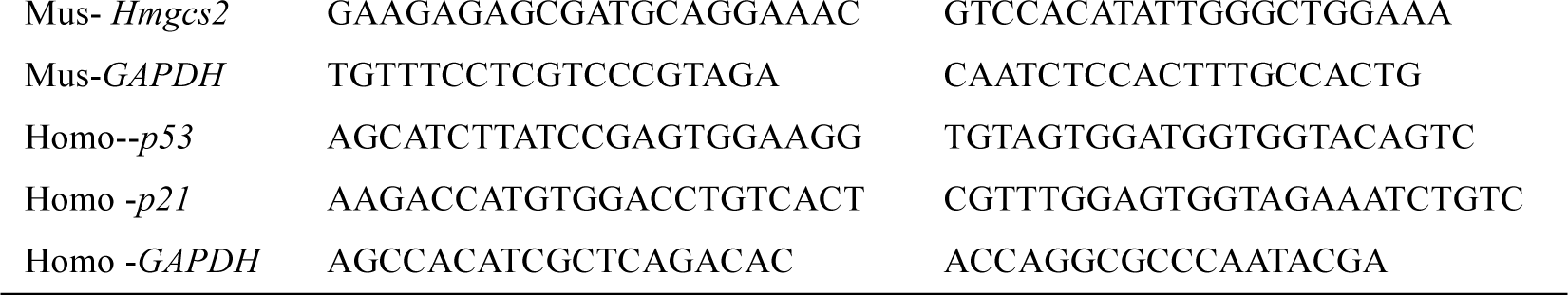
Primer sequences for qPCR analysis (qRT PCR)

**Fig. S1.**
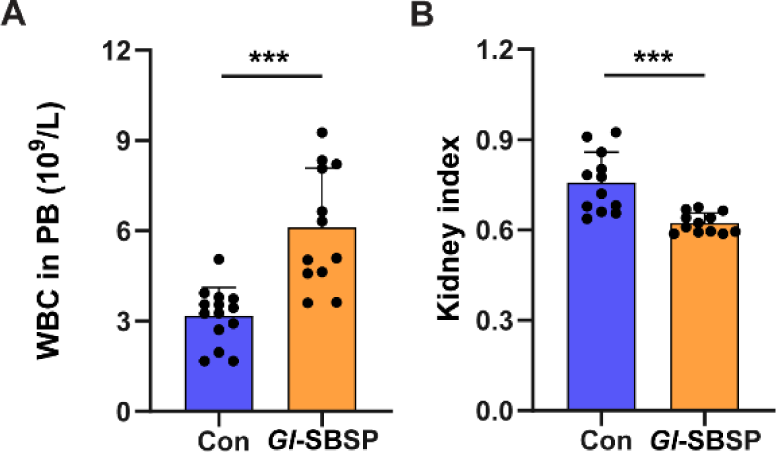
***Gl*-SBSP alleviates kidney aging in aged mice**. (A) Evaluation of WBC counts in PB. (B) The Kidney index of aged mice. * *P* < 0.05, ** *P* < 0.01, *** *P* < 0.001, Student’s *t*-test.

**Fig. S2.**
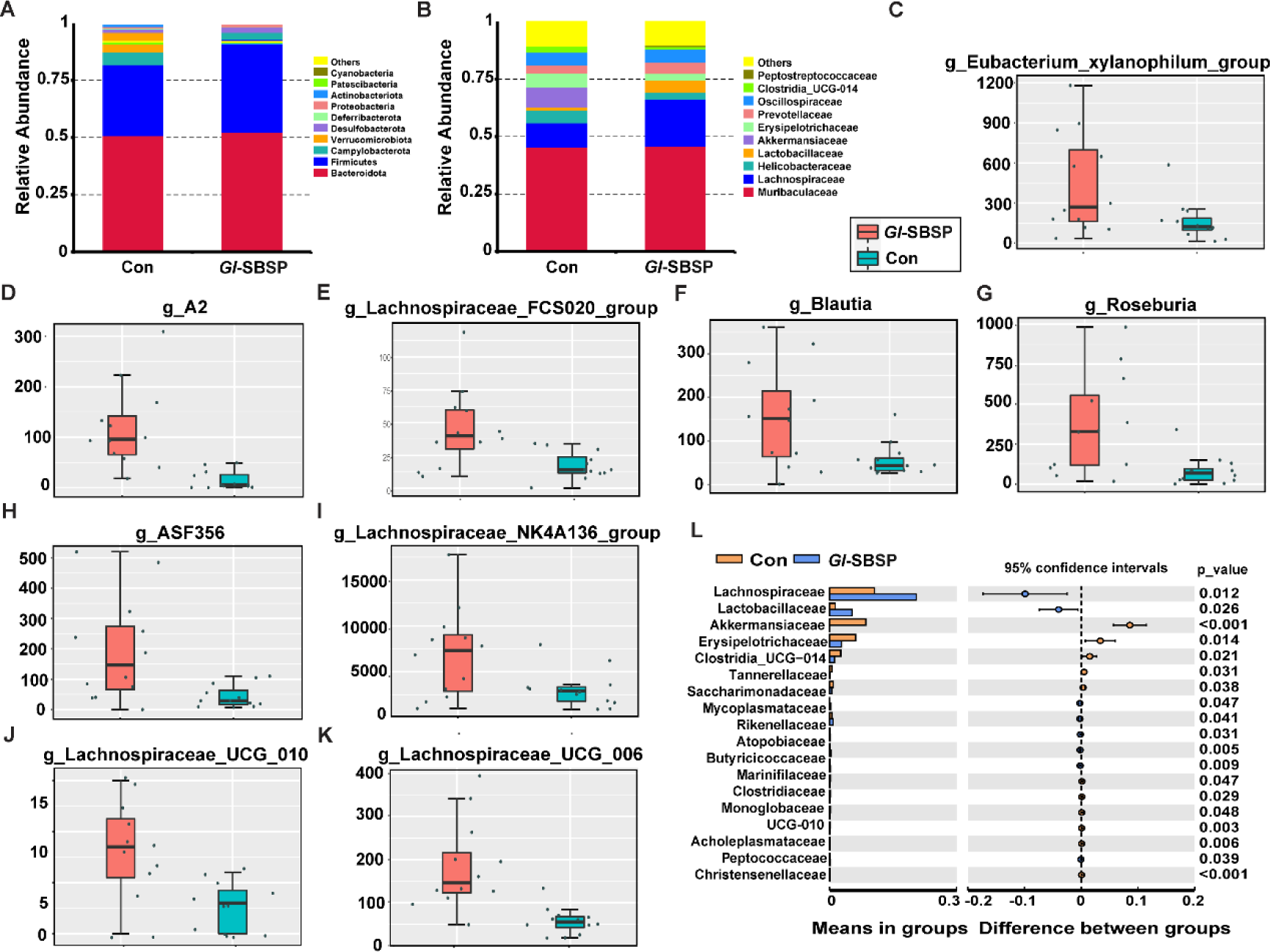
***Gl*-SBSP modulation of gut microbiota affects kidney aging in aged mice**. (A) Relative abundance of the top 10 abundant bacteria at the phylum level. (B) Relative abundance of the top 10 abundant bacteria at the family level. (C-K) The abundances of the most varied strain bacteria at the genus level of Lachnospiraceae were assessed using 16S rRNA high-throughput sequencing. (L) T-test analyzing the difference in composition of gut bacteria.\**P* < 0.05, \*\**P* < 0.01, \*\*\**P* < 0.001 Student’s *t*-test.

**Fig. S3.**
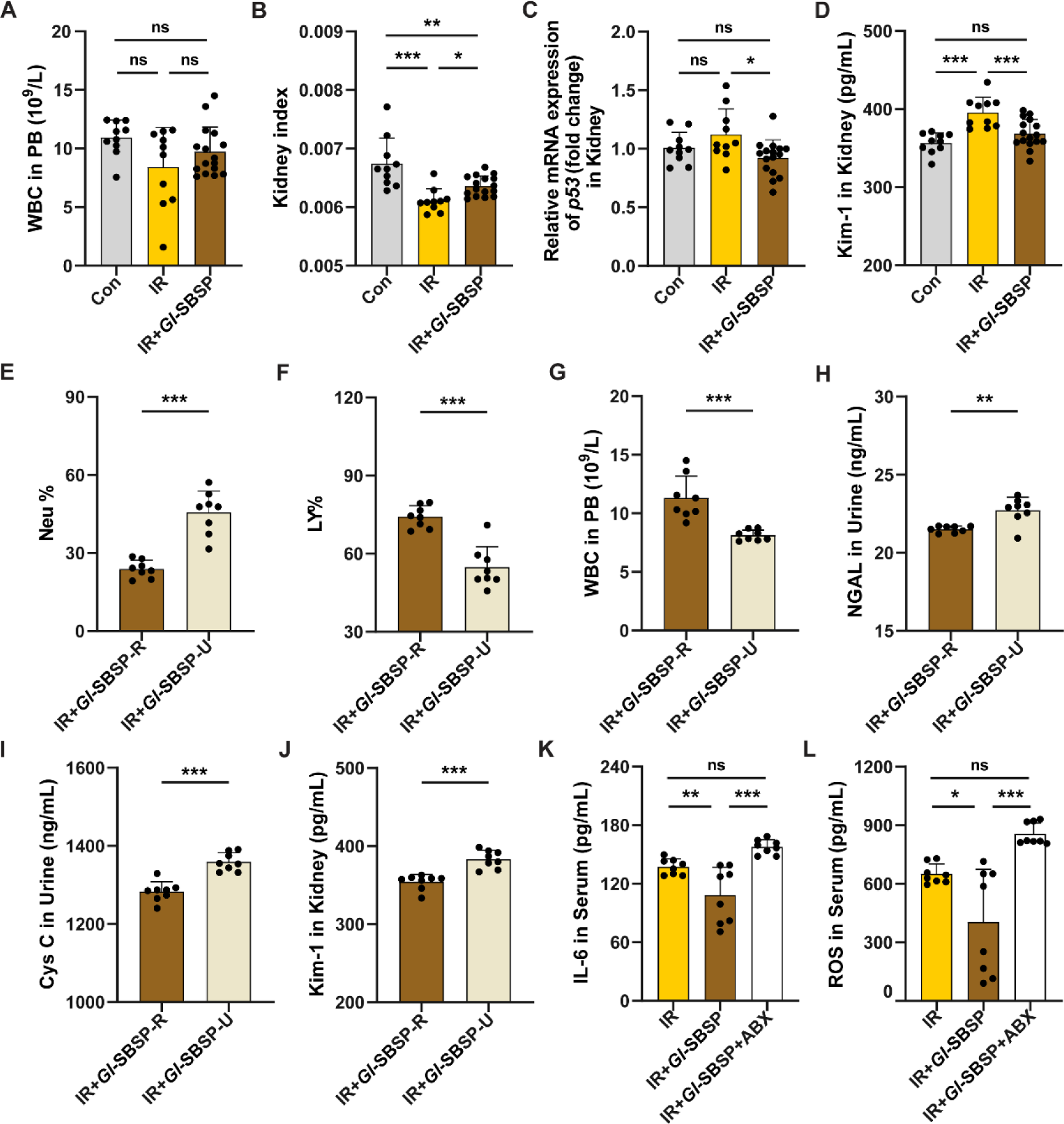
***Gl*-SBSP alleviates irradiation-induced kidney aging**. (A) Assessment of WBC counts in PB. (B) The Kidney index of mice. (C) The expression levels of *p53* in kidney tissues of mice. (D) Levels of Kim-1 in the urine of mice. (E-G) Percentage of Neu (E) and Lym (F) and WBC (G) counts in PB. (H-J) Levels of NGAL (H) and Cys C (I) and Kim-1 (J) in the urine of mice. (K-L) Levels of IL-6 (K) and ROS (L) in the serum of irradiated mice with ABX treatment. * *P* < 0.05, ** *P* < 0.01, *** *P* < 0.001, Student’s *t*-test between each two group, ANOVA among three groups.

**Fig. S4.**
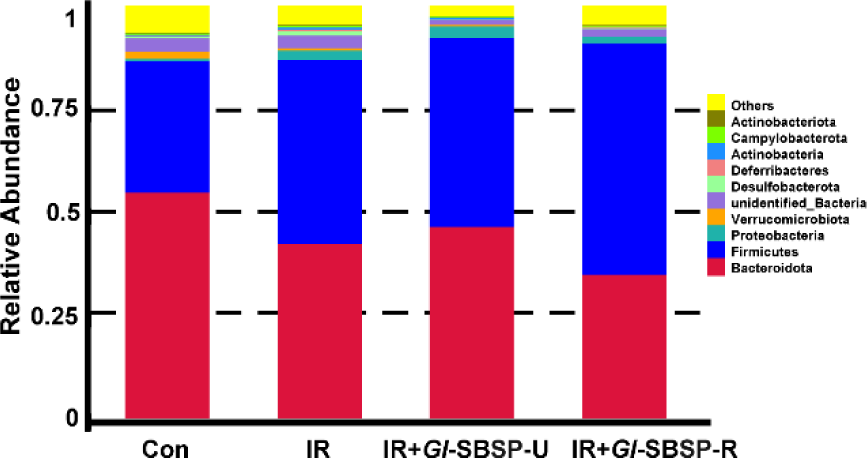
Relative abundance of the top 10 abundant bacteria at the phylum level in four groups.

**Fig. S5.**
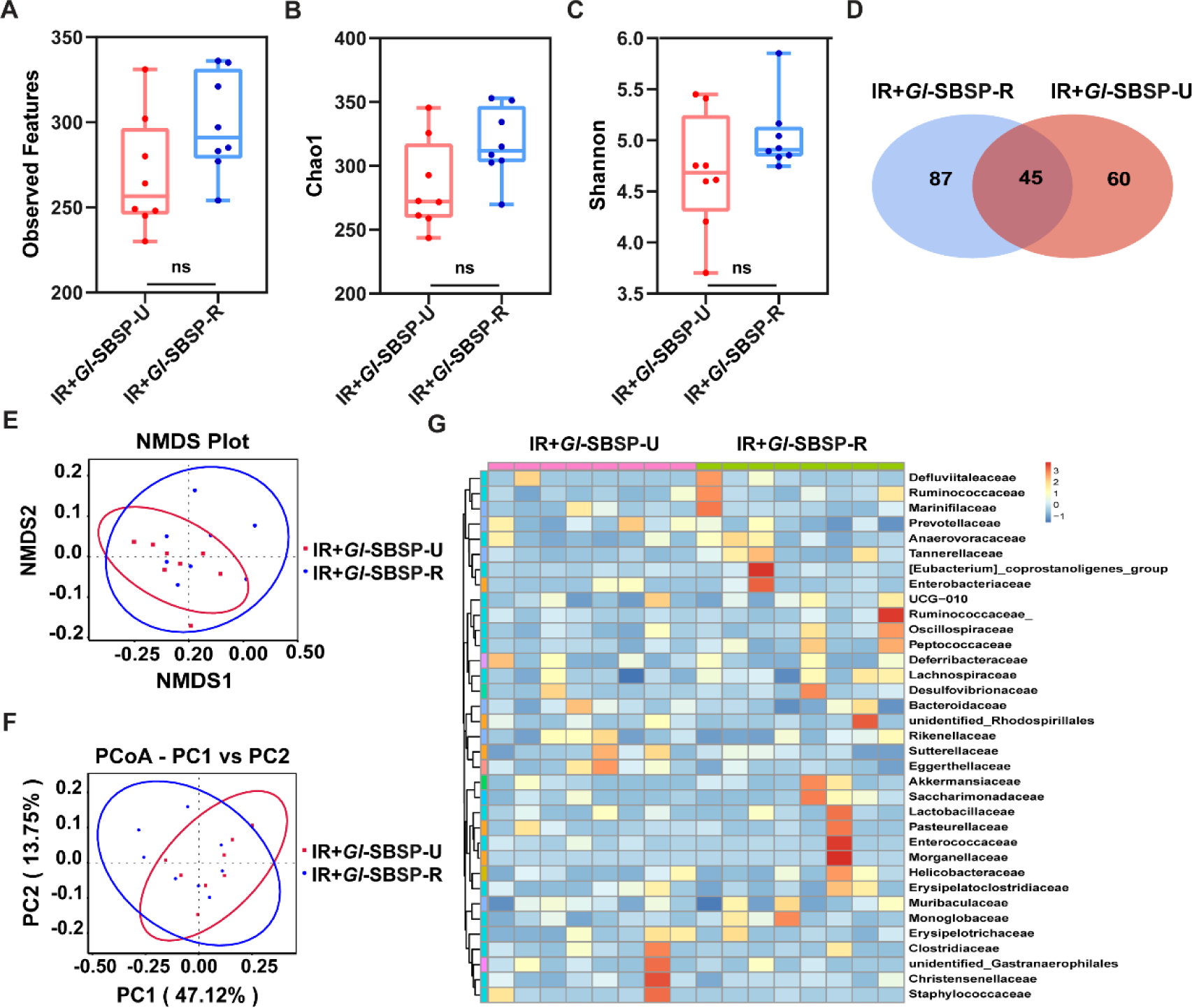
Intestinal bacterial composition before irradiation. n=8 per group. (A-C) The Observed Features (A), Chao1 (B) and Shannon (C) diversity index of intestinal bacteria were examined by 16S rRNA high-throughput sequencing. (D) The Venn diagrams illustrating the intersections and differences among the four groups. (E-F) NMDS (E) and PCoA (F) were used to measure the shift in intestinal bacterial composition profile. (G) The alteration of intestinal bacterial patterns at the genus levels. The heatmap is color-coded based on row Z-scores, with the highest and lowest bacterial levels shown in red and blue, respectively.

**Fig. S6.**
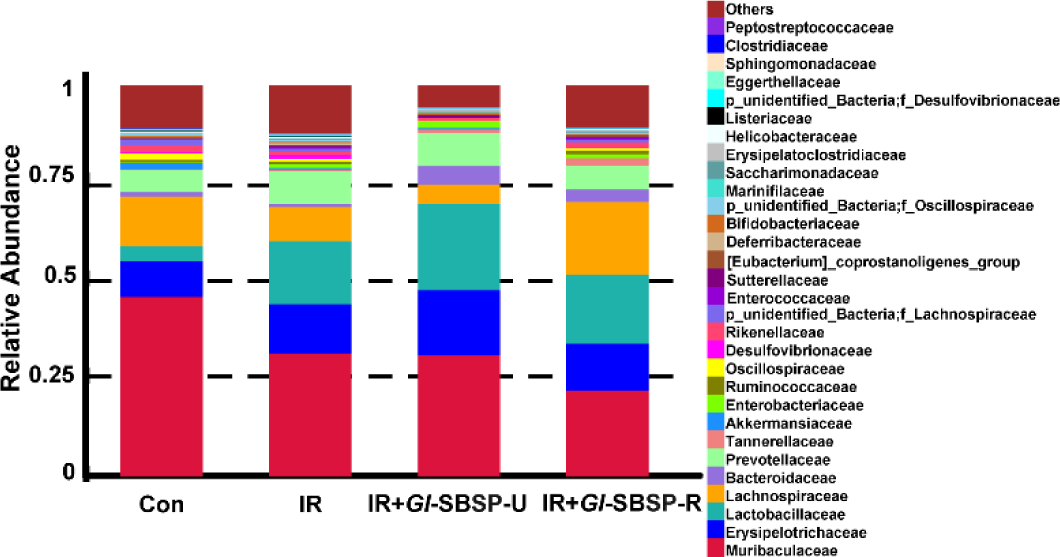
Relative abundance of the top 30 abundant bacteria at the family level in four groups.

**Fig. S7.**
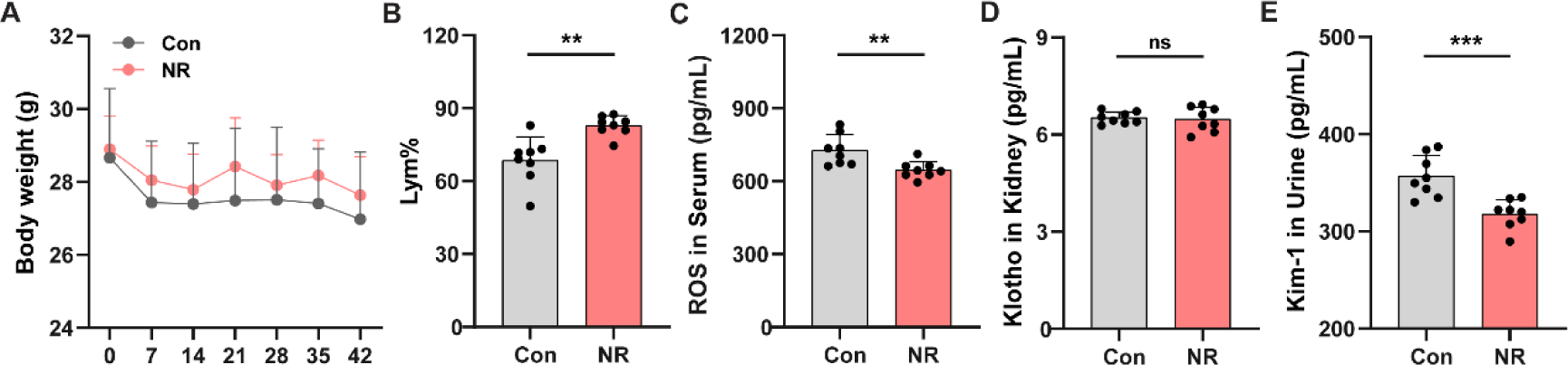
***Gl*-SBSP alleviates kidney aging through elevating NR in naturally aged mice**. (A) The body weight of aged mice. (B) Percentage of Lym in PB of aged mice was examined at 6 weeks. (C) Levels of ROS in the serum of aged mice. (D) Levels of Klotho in kidney tissues of aged mice. (E) Levels of Kim-1 in the urine of aged mice. * *P* < 0.05, ** *P* < 0.01, *** *P* < 0.001, Student’s *t*-test.

**Fig. S8.**
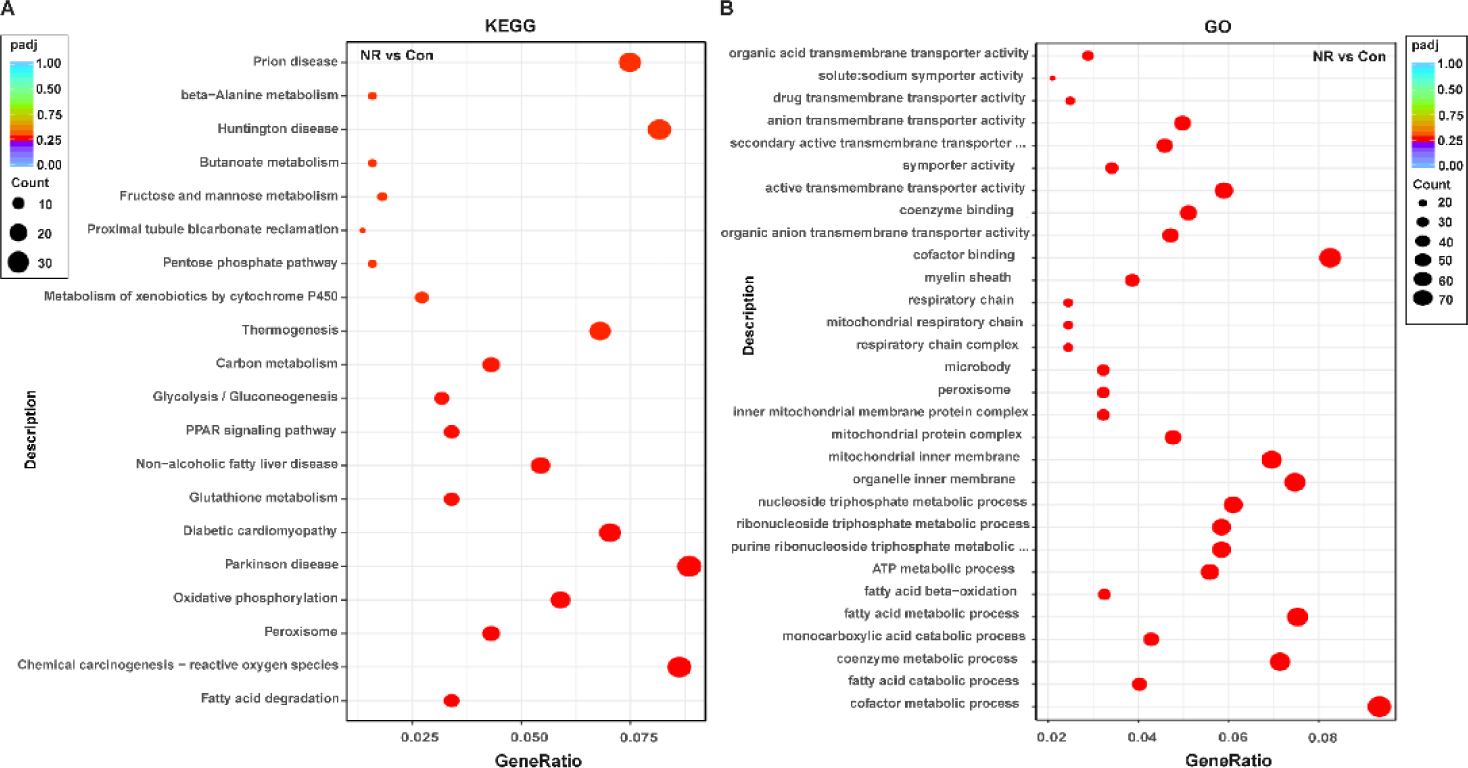
**NR relieved kidney aging via via regulating lipid metabolism**. (A) KEGG pathway analysis of the upregulation of differential metabolites in two groups. (B) GO pathway analysis of the upregulation of enriched metabolites in two groups.

**Fig. S9.**
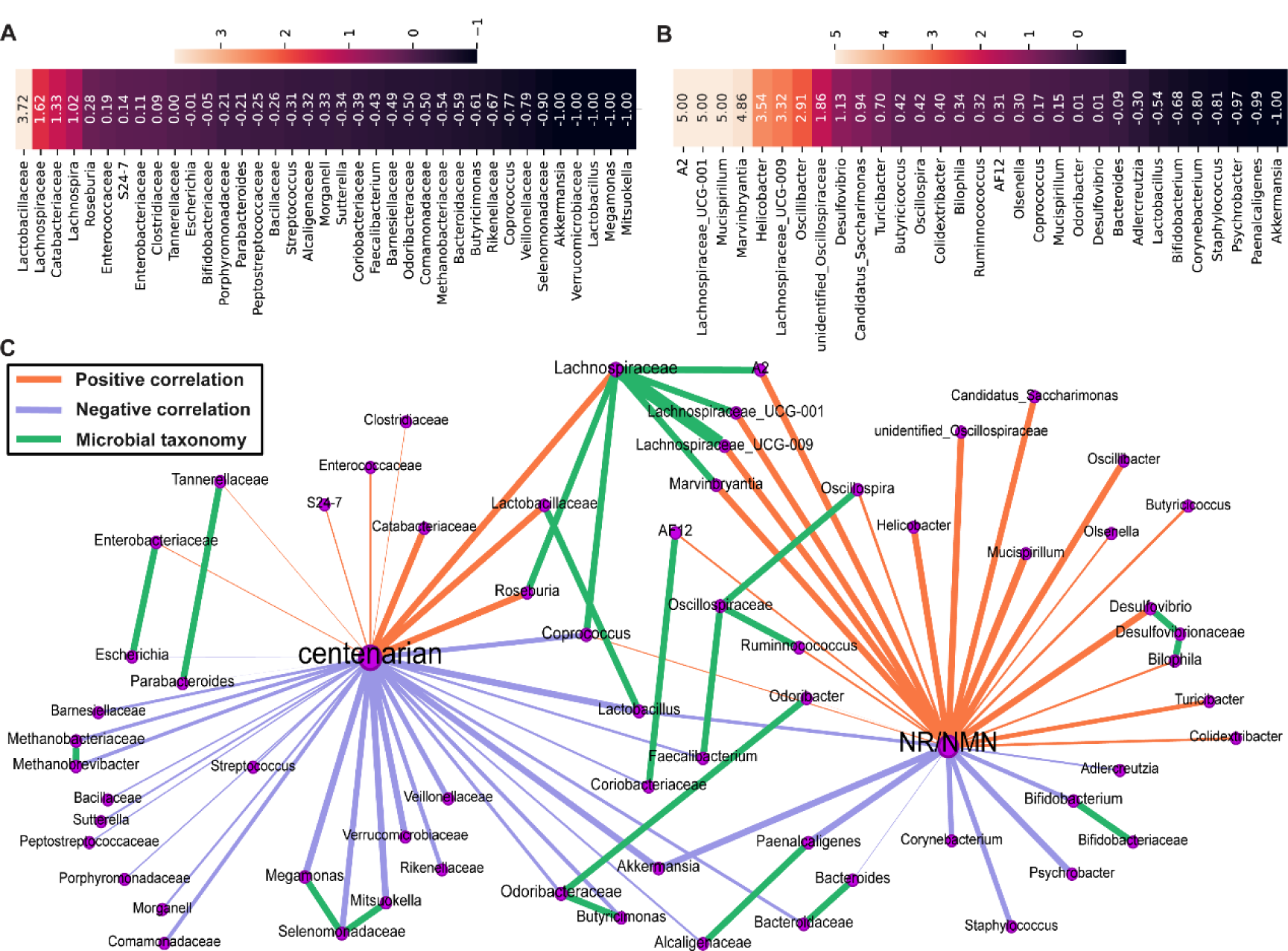
**Correlation analysis between Lachnospiraceae and NR**. (A) At the family level, the growth rate of gut bacteria in centenarians compared to the elderly. (B) At the genus level, the growth rate of gut bacteria in NR treatment compared to the control group. (C) Network graph of the association between gut microbiota in centenarians and NR/NMN-treated mice.

**Fig. S10.**
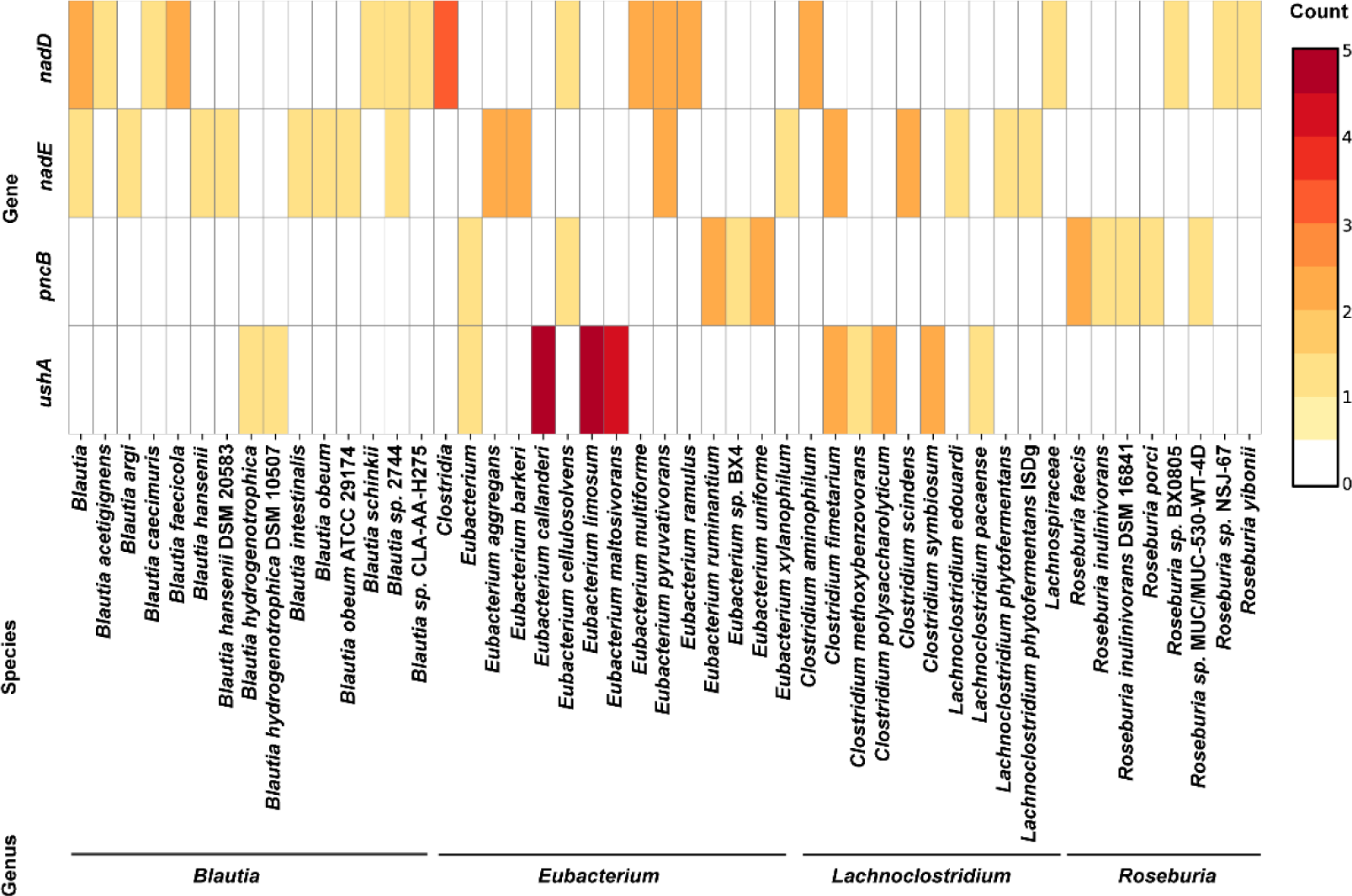
NR synthesis genes in Lachnospiraceae genome.

